# Dissection of *Plasmodium falciparum* developmental stages with multiple imaging methods

**DOI:** 10.1101/2020.06.25.171231

**Authors:** Katharina Preißinger, Beáta Vértessy, István Kézsmárki, Miklós Kellermayer

## Abstract

Efficient malaria treatment is a global challenge, requiring in-depth view into the maturation of malaria parasites during the intraerythrocytic cycle. Exploring structural and functional variations of the parasites through the intraerythrocytic stages and their impact on red blood cell (RBCs) is a cornerstone of antimalarial drug development. In order to trace such changes in fine steps of parasite development, we performed an imaging study of RBCs infected by *Plasmodium falciparum*, using atomic force microscopy (AFM) and total internal reflection fluorescence microscopy (TIRF), further supplemented with bright field microscopy for the direct assignment of the stages. This multifaceted imaging approach allows to reveal structure–functionality relations via correlations of the parasite maturation with morphological and fluorescence properties of the stages. We established identification patterns characteristic to the different parasite stages based on the height profile of infected RBCs, as obtained by AFM, which show close correlation with typical fluorescence (TIRF) maps of RBCs. Furthermore, we found that hemozoin crystals exhibit a strong optical contrast by quenching fluorescence. We demonstrate that these topographic and optical features also provide a tool to locate the hemozoin crystals within the RBCs and, in turn, to follow their growth.

---

Every year, more than 200 million people are infected with malaria. Five species of the *Plasmodium* genus cause human malaria infection, among which *P. falciparum* and *P. vivax* are the most widespread and mainly responsible for severe malaria [1]. The protozoon is transmitted into the human body by mosquito bite. Following the liver stage, an asexual cycle of the parasites takes place in the blood stream: They invade RBCs as merozoites and mature to rings, then to trophozoites and finally to schizonts when they multiply, burst out of the old host cells and start the next cycle by invading new RBCs. This intraerythrocytic cycle has been the subject of intense research because it is the cause of clinical symptoms and therefore the major target of antimalarial treatment and diagnosis. The digestion of hemoglobin by all *Plasmodium* species results in the accumulation of a micro-crystalline metabolic byproduct, the hemozoin, and in morphological changes of the RBC, which are typically characterized with bright-field microscopy (BFM) [2,3]. The development of a cavity associated with early-ring forms were observed within the RBCs, which spreads and extends for late rings. Towards the trophozoite stage, when a considerable portion of the hemoglobin is already transformed to hemozoin, the hemozoin becomes concentrated in a single food vacuole of the parasites in form of co-aligned crystals. During the trophozoite stage, the vacuole is settled to the side of the parasite, while it still grows in size. The prominent cavity disappears during maturation to the schizont stage[4]. These changes are likely associated with alterations in the external morphology and mechanical properties of the RBCs, which are little understood.

In order to explore *Plasmodium*-induced morphological and mechanical changes of infected RBCs, we combined AFM [5–9] and TIRF [10]. This comparative analysis, carried out on a large number of RBCs containing ring, trophozoite and schizont forms, allowed to correlate the morphological changes of RBCs with the *P. falciparum* developmental stages, which were identified by BFM. We could establish typical RBC height profiles, which are characteristic to the different stages. This provides the basis for an AFM-based identification of the parasites stages without the need of contrast materials or biochemical processes, which is a clear advantage with respect to most of the methods commonly used for the classification of malaria developmental stages, such as BFM microscopy on Giemsa stained smears [11], Polymerase Chain Reaction (PCR)[12] and flow cytometry [13]. In addition, we found that TIRF can be used to locate the hemozoin crystals within the RBCs, again in a label-free manner. We are going to extend these studies, carried out on fixed cells, by investigating infected RBCs with AFM and TIRF in cold aqueous solutions, slowing down parasite maturation. This protocol may provide conditions close to those realised in the human body to trace morphological and mechanical properties of the RBCs *in situ* during the maturation of the parasites.

## RESULTS

### Morphological changes in infected RBCs

A systematic AFM study was performed on various Giemsa stained thin film smears of synchronized *P. falciparum* cultures, i.e. individual smears contained mostly parasites of a certain developmental stage. The developmental stages —ring, trophozoite and schizont— were determined by BFM, as shown by representative images in the first row of Fig. 1. Co-imaging of the smears by the two methods did not facilitate the use of immersion objectives and, thus, compromised the resolution of the BFM images. In spite of the lower resolution, characteristic features of the stages are still well discernable in the BFM pictures. Healthy RBCs show almost no surface deflection and appear as nearly flat disk in their partially dehydrated form within the smear, as shown in the AFM image of Fig. 1a. There is an abrupt change in the ring stage, where a cavity develops at the position of the ring, typically close to the center of infected RBC. This is a hallmark of RBCs infected by ring-stage parasites, which is clearly resolved in the corresponding topography image in Fig. 1b. With maturation to the trophozoite stage, the indentation widens, extending up to typically half of the cell diameter, and gets off-centered (see Fig. 1c). The topography in the schizont stage, displayed in Fig. 1d, reflects another radical transformation. The surface of the RBC becomes flat and featureless again, similarly to the ring stage, though it shows a strong roughness. In addition, the contour of the cell is less regular, in contrast to the nearly cylindrical shape of the uninfected RBC and the early stages. These may indicate the extension of the parasite cytoskeleton over the RBC.

**FIG. 1.**
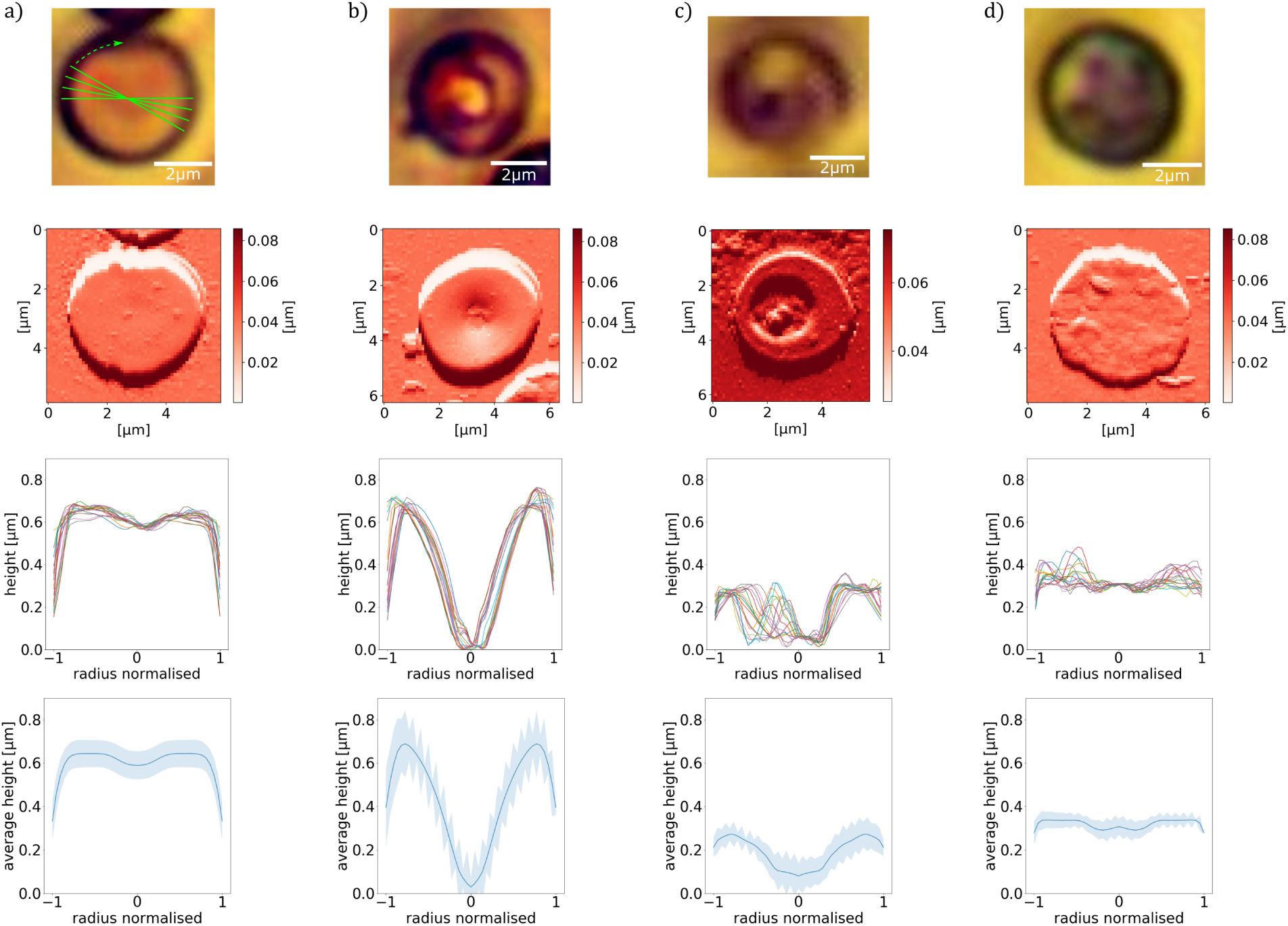
Morphology of RBCs infected by P. falciparum. Results on RBCs with no parasite, with ring, trophozoite and schizont stages are shown in columns (a), (b), (c) and (d), respectively. First row: BFM images of RBCs in stained smears. Second row: AFM topography images recorded over the same areas as the BFM images above. Third row: Radial height profiles of RBCs as obtained by radial cross-sections of the AFM topographic images above. Green lines in the top-left panel indicate the path of the radial cross-sections. Forth row: Mean height profiles obtained by averaging the different cross-sections shown in the panels above. The standard deviation of the height is indicated by the shaded region around the curves.

Next, we studied if the morphological information can be extracted from the 2D images in the form of 1D height profiles, while preserving the typical features of the different stages. For this purpose, we developed a python program to determine the height profile along an arbitrary number of cross-sections through the center of the cell. In the present study, we selected 18 radial cross-sections, i.e. made a cut in every 10*°*, as indicated schematically in the top panel of Fig. 1a. The height profiles along these radial cuts, as obtained for the exemplary AFM images in the second row of Fig. 1, are shown in the third row of Fig. 1 for each parasite stage. All the height profiles, apart from those of the trophozoite stage, are nearly symmetric to the centre of the cell. It is also clear that the height shows small variations along the radius for uninfected cells and schizonts, while it exhibit a large drop in rings and trophozoites, when approaching the cell center from the periphery. Averaging these height profiles for a given cell reveals the radial dependence of the height, as seen in the forth row of Fig. 1.

In the average height profiles of the different stages we can recognize the typical features already noticed in the 2D images: The flat discoid shape of the uninfected RBC [14], the central cavity signaling the ring stage [4] and the nearly featureless landscape with reduced overall height, characteristic to the schizont stage. However, by this averaging we loose the information about the asymmetry of the cells, arising e.g. from the off-centering of the parasites, which is particularly important for trophozoites. This information can be recovered from the variance of the hight, as represented by the shaded area around the average curve. This variance is not solely determined by the overall asymmetry of the height profiles, but short-scale surface roughness can also contribute to it. However, the observations, that the variance is the largest for the trophozoite stage and increases gradually from the periphery to the center, imply the primary role of asymmetry in the variance of the height. In the following, we will show that these morphological features are not specific to the exemplary cells displayed in Fig. 1, but characteristic to the different stages.

For further analysis, we sorted all topography images according to the parasite stages into the four categories —uninfected RBC, ring, trophozoite, schizont—, using BFM. In order to identify key differences between the different stages, the radial height profiles were determined for all individual cells by the averaging process described above. Then, these radial height profiles were averaged for all cells of the same stage (approx. 100 cells for each stage) to obtain a master curve describing the typical morphology of that stage. These master curves, shown in Fig. 2, reproduce the key features observed for individual cells of the corresponding stage (see Fig. 1).

**FIG. 2.**
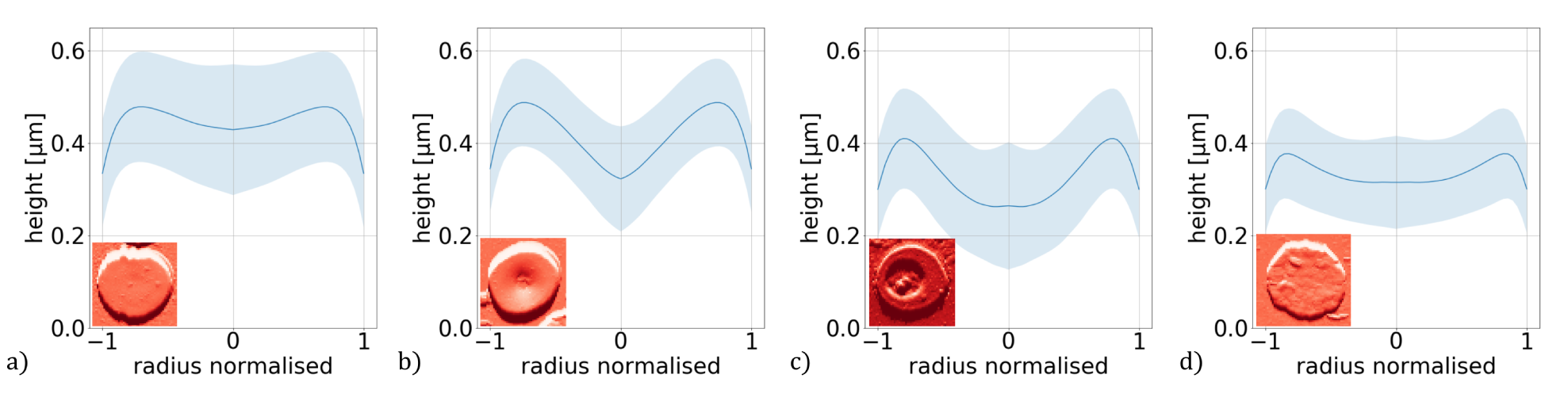
Master curves representing the typical height profiles of RBCs containing no parasites (a) and ring (b), trophozoite (c) and schizont (d) stages of *P. falciparum*. The height profile curve typical for a given stage were obtained by averaging the radial height profile curves of approx. 100 individual RBCs corresponding to the same stage. The dark region indicate the standard deviation. The radial height profile curves for individual RBCs are shown in the last row of Fig. 1.

The mean height is nearly the same for the uninfected RBC and for the ring stage, while it shows a significant reduction in the trophozoite and schizont stages. More interestingly, the height drop from the maximum at the cell edge to the minimum near the center shows a more pronounced dependence on the stages. The change in the height along the radius is ∼10%, ∼35%, ∼45% and ∼20% of the mean height for the uninfected RBC and for the ring, trophozoite and schizont stages, respectively. These values together with values of the mean height are also displayed in Fig 3. In addition, the variance of the height is indicated as a shaded area around the master curves, which shows an increase towards the center for the trophozoite stage, while its radial dependence is weaker in the other three cases. Our observations show that these are valuable parameters to identify the developmental stages, comprising an essential part of the information encoded in the 2D images.

**FIG. 3.**
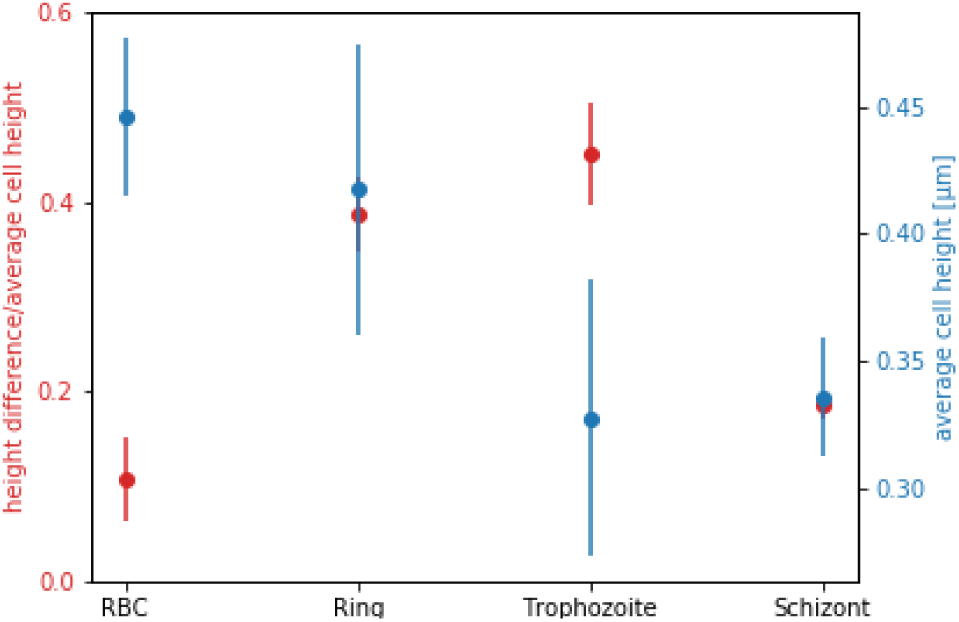
Mean height of the RBC and difference between maximum and minimum height divided by mean height along the radius for different developmental stages of *P. falciparum*. These parameters were calculated from master curve of each stage, shown in Fig. 2.

### Fluorescence properties of *P. falciparum*

The maturing of malaria parasites not only changes the morphology, but also the optical properties of the infected RBC. Notably, our TIRF study, carried out with an excitation wavelength of 405 nm, reveals the emergence of autofluorescence in dried smears. This wavelength was chosen to be in resonance with the strongest absorption peak in protoporphyrin IX, which plays an important part in the biosynthesis of heme[15,16]. Our measurement samples were parasite enriched smears on cover slips, which were prepared from cultures treated with Percoll (see). Simultaneously we recorded BFM images on the same samples to identify the stage of the parasites. Based on the recorded fluorescence intensity profile, we performed a similar evaluation as for the topography in the AFM images. We cut the cell into radial cross-sections and plot the intensity of the TIRF images along these radial cuts. Figure 4 shows a TIRF image representative for each stage with the corresponding 1D intensity profiles.

**FIG. 4.**
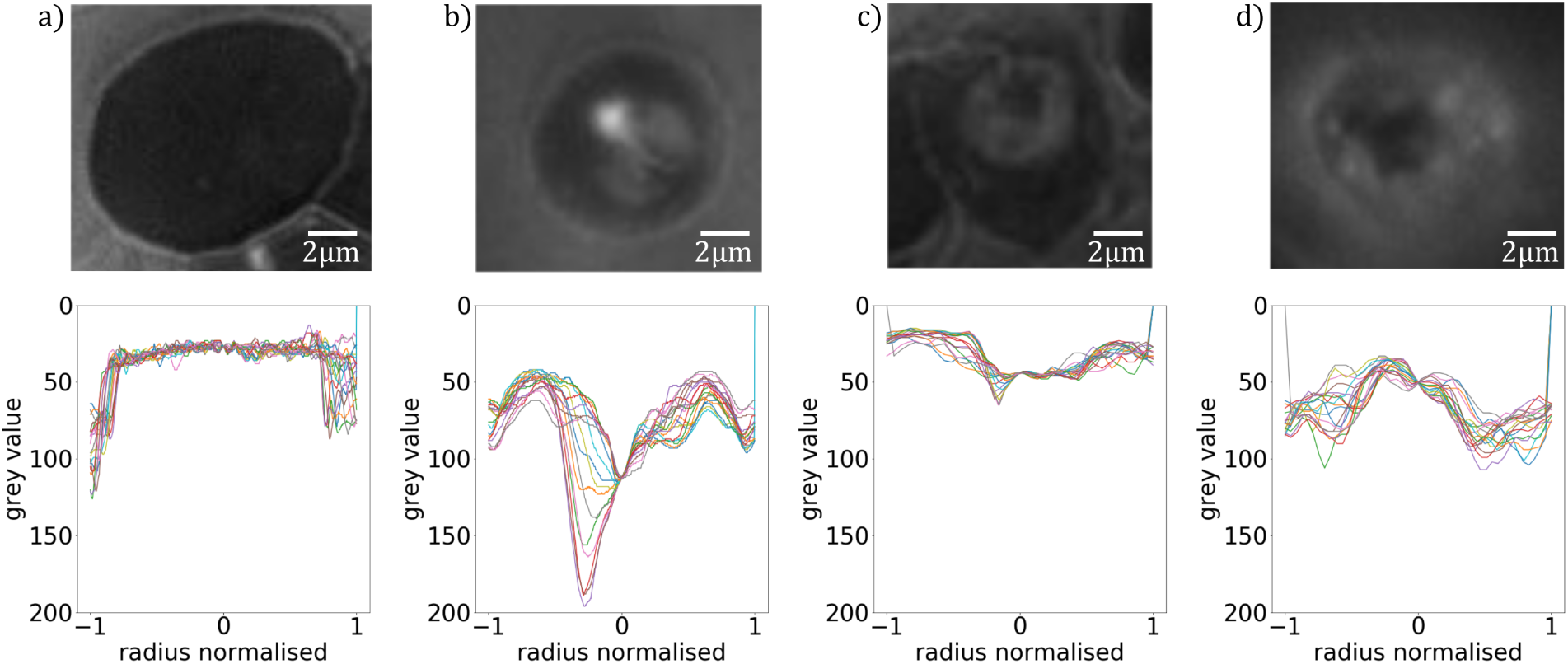
Fluorescence properties of RBC with no parasite (a), with ring (b), with trophozoite (c) and schizont (d) stage *P. falciparum* parasites, as studied by TIRF. First row: TIRF images of RBCs in thin smears. Second row: Radial fluorescence intensity profiles for the different stages, obtained as radial cross-sections of the images above.

Since TIRF is a surface sensitive method with a penetration depth of approx. 100 nm[17], the images are not representative for the whole cell depth, only for the region close to the cell-glass interface. For an easier comparison with the AFM results, the spectra were turned upside down, i.e. the intensity increases from the top to the bottom. The uninfected RBC shows no fluorescence, as evident from the low and uniform signal in Fig. 4a, detected over the whole area of the cell. In the ring stage, small spot with high fluorescence intensity is observed, which is connected to a more extended region of moderate fluorescence, embedded in the otherwise non-fluorescent cell (Fig. 4b). An central region in trophozoite stage emits a diffuse signal surrounded by a non-fluorescent edge region of the cell (Fig. 4c). The diffuse region extends nearly over the cell diameter in the schizont stage, but the quenching of the fluorescence is found close to the center, the dark spot in the middle of Fig. 4d. In fact, a similar black region, though smaller, can also be observed for the trophozoite stage. The reason for the quenching of fluorescence will be discussed later.

## DISCUSSION

By combining AFM, TIRF and BFM, we characterized the developmental stages of *P. falciparum* parasites using in thin smears of synchronized cultures. Our data demonstrates that the maturing of the parasite not only changes the morphology of the RBC but also activates fluorescence. With a statistical analysis of the height profiles of the cells, we could derive an identification pattern for each developmental stage. The average height together with the maximum height difference and the variance of the master curves clearly characterises the stages. Our results on morphology support the observations reported about the maturation of malaria parasites by Grüring *et al*. [4]. While the uninfected RBC, the ring and schizont stages are rather symmetrical to the center of the cell, the symmetry is often disrupted in the trophozoite stage. Grüring *et al*. showed that the food vacuole settles to the side of the trophozoite, which might explain the asymmetric behaviour of the height profiles in Fig. 5a. With maturation to the schizont stage, they observed the shifting of the food vacuole back to a more central position [4]. Therefore, the central peak in the height profile of schizonts, as shown in Fig. 5b, is likely to be the food vacuole.

**FIG. 5.**
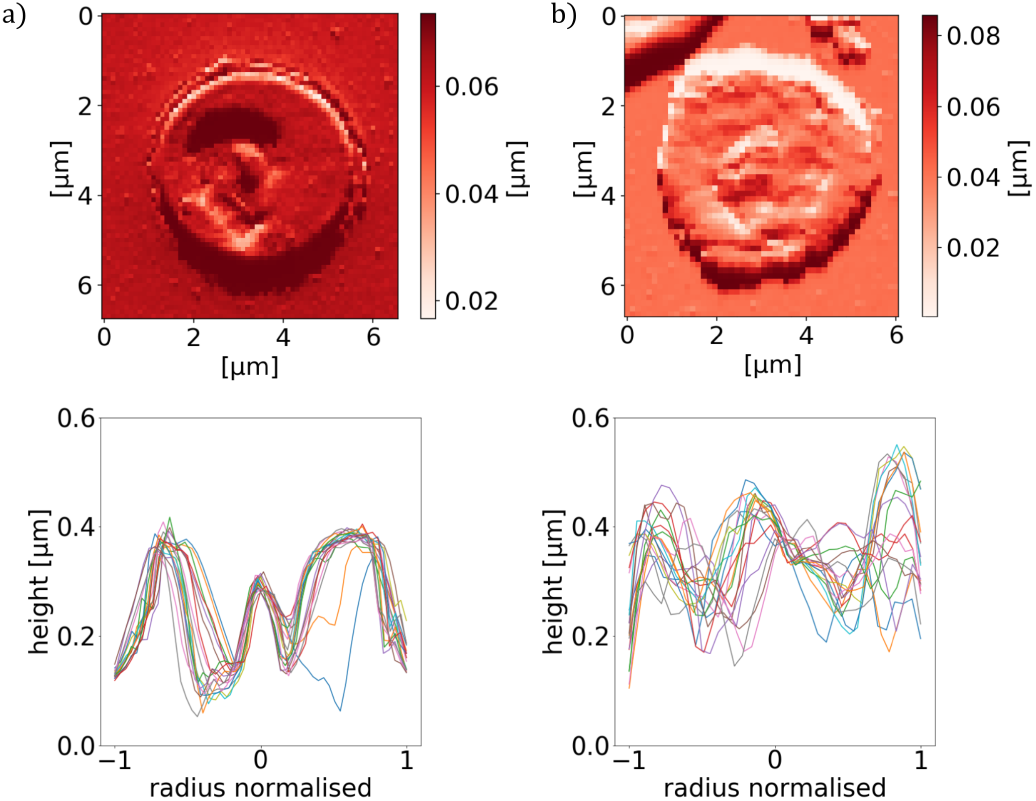
AFM images (top panels) and the corresponding radial cross-sections of the height (bottom panels) for an RBC containing a trophozoite/schizont in coloumn (a)/(b). The cross-section in both cases shows a maximum around the center of the cell.

The combined BF and TIRF imaging provided new information about the autofluorescence properties of malaria parasites. We could show that uninfected cells emit no fluorescence, while the parasite becomes fluorescent during maturation. The signal changed from a well defined indentation in the ring stage to a diffuse pattern in the schizont stage. In the signal of trophozoite (Fig. 6a and b, middle) and schizont stage (Fig. 6a and b, right), we identified non-fluorescent regions inside the fluorescent parasite. An additional TIRF study on extracted hemozoin crystals showed no fluorescence emission (Fig. 6a and b, left). This highly suggests that we can image malaria pigment with TIRF. However, it is important to note that we only see up to 100 nm deep into the parasite. Such signatures of the food vacuole can indeed be resolved in the other exemplary AFM images of trophozoite and schizont stages in Fig. 1c & d, though less pronouncedly in the schizont stage.

**FIG. 6.**
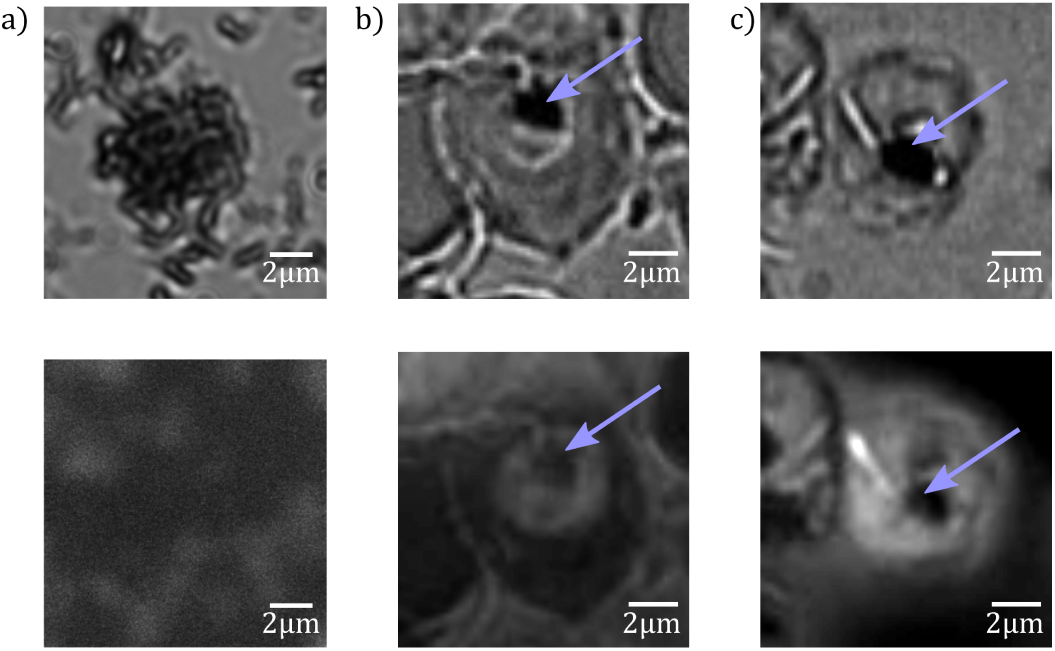
BFM (top panels) and TIRF (bottom panels) images of extracted hemozoin crystals in column (a) and of an RBC containing a trophozoite and schizont in column (b) and (c), respectively. The extracted hemozoin crystals show no fluorescence. In the trophozoite and schizont, the non-fluorescent regions are indicated by blue arrows.

When comparing the fluorescence maps with the topography images, respectively shown in the bottom line of Fig. 4 and the third row of Fig. 1, we observe a clear correlation: The features characteristic to the different parasite stages appear in the two types of images in a similar fashion. Most interestingly, our hypothesis that the central peak observed in the height profiles for trophozoites and schizonts corresponds to the food vacuole is supported by TIRF studies on the same stages. Figs. 6b & c display BFM and TIRF images simultaneously recorded on RBCs containing trophozoites and schizonts, respectively. The food vacuole appears as a dark spot within the parasite in both cases. In the latter case, the darkness is due to the quenching of fluorescence by the hemozoin crystals. The non-fluorescent nature of hemozoin is evidenced in Figs. 6a by TIRF on extracted hemozoin crystals.

Our experiments clearly illustrate the connection between RBC morphology, fluorescence properties and the maturing of *P. falciparum* and these observations can help to establish structure–functionality relations, to reveal the detailed mechanism governing the intracellular parasite activity and eventually to guide the development of new antimalarials. Nevertheless, the present study, performed on fixed cells, has some inherent limitations. Under such conditions the shape of the RBCs is deformed, i.e. the cells are flattened, and their mechanical properties are also altered by partial dehydration. Furthermore, the smears only show a snapshot of the parasite life cycle. By extending the experiments to living parasites, more detailed and realistic information about the parasite maturation can be obtained. However, performing multi-probe imaging under conditions close to those in the human body requires simultaneous liquid AFM and TIRF studies on parasite cultures, which is a highly challenging task.

## MATERIALS AND METHODS

### Sample preparation

*Plasmodium falciparum* parasites from the laboratory adapted strain 3D7 were cultured in culture medium (Albumax, 25 mg/L Gentamycin, RPMI 1640) and maintained in an atmosphere of 5 % CO_2_ and 5 % O_2_[18]. The cultures were raised to 5 % parasitemia and treated with Percoll or Sorbitol to synchronise the parasites. For the AFM measurements, we sorted the schizont stage parasites from other stages. For this, cultured parasites were added to 5 ml of Percoll and centrifuged, layering them according to their densities. The schizonts, having the smallest density of the developmental stages, could be recovered from the first layer[19]. After centrifugation, 1.2 µl of the remaining pellet of RBC with enriched parasitemia were used to make a thin film smear on a glass slide. This procedure was repeated for ring stage parasites, which were sorted by Sorbitol treatment[10]. To classify the parasites prior to the morphology measurement, the smears were stained with Giemsa’s stain[11].

### Morphological measurements

For the morphological measurements of the smears, we used a MFP-3D AFM from Oxford Instruments. It was operated in AC mode at a scanning speed of 0.25 Hz. Scan areas of 90 × 90 µm were conducted to analyse the morphology of RBC and iRBC. Each scan was performed with a resolution of 512 × 512 pixels.

### Fluorescence measurements

For the analysis of fluorescence properties of *Plasmodium falciparum*, we used a TIRF microscope, which was operated at a wavelength of 405 nm. The smears were prepared on a cover slip and dried in air before the measurement.

### Evaluation program

#### Data acquisition

The data, we analysed was obtained from the height trace and the amplitude of each AFM scan, which were converted to txt files.

#### Preprocessing of image

The goal of the preprocessing was to detect each cell in the AFM image. For this, the txt file was converted to a binary image by thresholding, which enabled the detection of almost every cell. The algorithm did not work on overlapping or folded cells, but this was not a problem because we only used non deformed cells for the evaluation.

#### Characterisation of cell by height profile

To characterise the cells by height profile, we determined 18 cross-sections through the height trace, which were rotated in the cell plane by an angle of 10*°*, so that every pixel along the outline of the cell was covered. From these, we determined an average height profile to describe a single cell by averaging the height of an arbitrary number of radial sections in the cell.

## ACKNOWLEDGMENTS

The research carried out here was supported by the National Reserach, Development and Innovation Office of Hungary (K119493, VEKOP-2.3.2-16-2017-00013 to BGV, NKP-2018-1.2.1-NKP-2018-00005), the BME-Biotechnology FIKP grant of EMMI (BME FIKP-BIO), the BME-Nanotechnology and Materials Science FIKP grant of EMMI (BME FIKP-NAT), the National Heart Programm (NVKP-16-1-2016-0017) and SE FIKP-Therapy Grant.

